# Ancient DNA of the Toronto Subway Deer Adds to the Extinction List of Ice Age Megafauna

**DOI:** 10.1101/2025.09.15.676284

**Authors:** Camille Kessler, Oliver Haddrath, Burton K. Lim, Aaron B. A. Shafer

**Affiliations:** Environmental and Life Sciences Graduate Program, Trent University, Peterborough, Ontario, Canada; Division of Evolutionary Biology, LMU Munich, Planegg-Martinsried, Bavaria, Germany; Department of Natural History, Royal Ontario Museum, Toronto, Ontario, Canada; Department of Forensic Science, Trent University, Peterborough, Ontario, Canada

**Keywords:** Quaternary, museomics, Great Lakes region, aDNA, Cervidae

## Abstract

The late Pleistocene was a time of global megafaunal extinctions that were particularly severe in North America. The continent lost many mammal taxa, but the validity of several remain ambiguous, including a high proportion of Cervidae taxa. *Torontoceros hypogaeus* is represented by a single specimen (ROMM75974) discovered in 1976 during excavation work for the Toronto subway in Canada. The species was described based on its unique antler morphology, but the variable nature of that trait and the species near absence in the fossil record leads to uncertainty concerning its systematic relationships. We used ancient DNA to clarify the taxonomic relationship and evolutionary history of *T. hypogaeus*. We performed mitochondrial and whole genome analyses with related cervids and showed that ROMM75974 has a close affinity, but relatively high divergence from the *Odocoileus* sister species. While some ambiguity remains, ROMM75974 could represent a distinct *Odocoileus* species to be included in the list of extinct North American taxa. This unique population was likely adapted to open landscape which was rapidly replaced with dense woodland in this region at the end of the Pleistocene, highlighting the role of climate change in the extinction of megafauna biodiversity at the end of the ice age.

## Introduction

The late Pleistocene was characterised by an extinction event that particularly affected large mammals[1,2]. The causes of these extinctions have largely been attributed to human impact and climate change, or a combination of the two[3–9], although other factors have been suggested[10,11]. This extinction event was severe in North America which lost between 33 and 37 large mammal genera[1–4,6,7,12,13], and while some genera are consistently included in summary lists of lost taxa (e.g., *Mammut, Equus*), others are not (e.g., *Holmesina, Capromeryx*). Four cervids (deer) disappeared during the Pleistocene: *Bretzia, Navahoceros, Torontoceros*, and *Cervalces*, only the latter two unequivocally went extinct during the late Pleistocene extinction event. Together with the extant *Odocoileus* and *Rangifer*, they represent the only genera in the deer family to have inhabited North America during the glacial cycles of the Pleistocene[14–17]. The extant wapiti (*Cervus canadensis*) and moose (*Alces alces*) originated in Eurasia and colonised North America through Beringia after the Last Glacial Maximum (LGM)[18–23].

Cervids first appeared in the North American fossil record approximately 5 million years ago (Mya) and consisted of three medium-sized genera: *Eocoileus, Bretzia*, and *Odocoileus*[14,24]. *Cervalces, Rangifer* and *Navahoceros* (American mountain deer) emerged later[14,25], although the taxonomic legitimacy of the latter is uncertain[26,27]. *Torontoceros hypogaeus* was described in 1982[28] and is represented by a single specimen (ROMM75974) dated at 11.3 thousand years ago (kya), noting this date is approximate as radiocarbon dating has drastically improved since 1982. The specimen consists of a partial cranium with incomplete antler beams, found in Toronto (Ontario, Canada) during excavation work for a metro line and was popularly dubbed the Subway deer. Similar in size to white-tailed deer (*Odocoileus virginianus*) and caribou (*Rangifer tarandus*), *T. hypogaeus* has a distinctive antler morphology that does not match any other American Cervidae[28] (Fig. 1, Fig. S1). Specifically, its antlers are heavy, with beams that are nearly horizontal without branches or palmation unlike caribou, and compared to *Odocoileus*, its brow tines are developed and asymmetrical[28].

**Figure 1:**
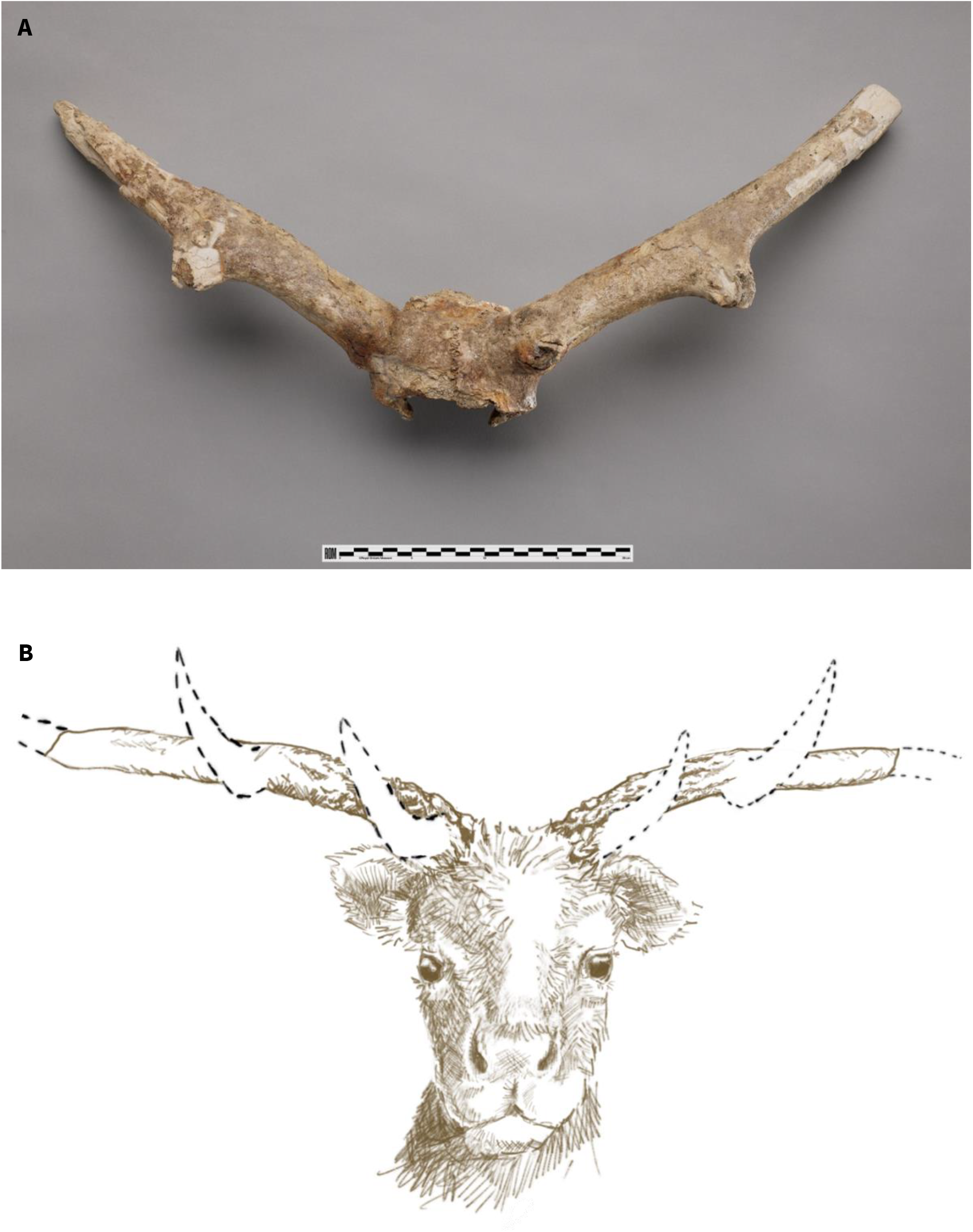
Specimen ROMM75974 of *Torontoceros hypogaeus* from the Royal Ontario Museum, Toronto. A) Photograph from anterior aspect, credit: Paul Eekhoff. B) Artistic rendering, credit: Sherri Owen

The genus *Torontoceros* was proposed primarily based on antler morphology, a highly variable and complex trait for which the use in systematics has been discouraged[14]. Two studies considered *T. hypogaeus* as a member of the genus *Rangifer*[22,29], either because the morphology resembled that of some North American caribou populations[29], or that it could reflect adaptation to open landscape[22]. The ambiguity around *T. hypogaeus* taxonomy is heightened by the rarity of this genus in the fossil record. Perhaps due to this uncertainty, *T. hypogaeus* has not been consistently included in analyses of the Pleistocene megafaunal extinction (Table S1). Here, we sought to clarify the taxonomic affinity of the Subway deer (*T. hypogaeus*) using ancient DNA sequencing. We carried out phylogenetic analyses of mitochondrial and whole genome sequencing data to resolve the taxonomic position of this enigmatic fossil within Cervidae, and to further clarify deer evolutionary history and extinction events in North America during the Pleistocene.

## Material and methods

### Ancient DNA extraction and sequencing

The specimen ROMM75974, described as the new genus and species *Torontoceros hypogaeus* by Churcher and Peterson[28], is deposited at the Royal Ontario Museum and consists of an incomplete cranium with partial antlers and bone fragments (Fig. 1, Fig. S1). We sampled a bone fragment from the posterior of the antler so as not to impact the integrity of the sample when exhibited. We performed aDNA extraction and library preparation in a dedicated laboratory at the Royal Ontario Museum. We irradiated the bone fragment with UV light for 11 minutes on all sides in a Stratagene UV Stratalinker 2400 before pulverizing it in liquid nitrogen with a mortar and pestle. We then extracted DNA from approximately 60 mg of bone powder following the silica column protocol from Dehasque et al. (2022)[30], and prepared double-stranded sequencing libraries with USER enzyme treatment following Meyer & Kircher (2010)[31] and including one extraction and one library prep negative control. Finally, we used unique barcodes to double-index and amplify the sequencing libraries as described in Diez-del-Molino et al. (2023)[32], with four independent PCR amplifications under the following protocol: 95°C for 2 min followed by 12 cycles of 95°C for 15 s, 60°C for 30 s and 68 °C for 1 min. We sent the amplified libraries to The Centre for Applied Genetics in Toronto, Canada, for paired-end sequencing of 150 bp reads on an Illumina Novaseq 6000 with SP flowcells.

For comparison with modern data, we collated 17 published mitochondrial genomes of Cervidae, and whole genome resequencing data from six deer species (Table S2, Table S3, Fig. S2), including all Northern American representatives (USA & Canada). Resequencing data was not available for all species investigated with mitochondrial genomes.

### Data processing

For the ROMM75974 sample data preparation, we trimmed and merged the reads in SeqPrep (v1.2)[33] with default settings and minor source code edits[34]. We mapped the merged reads to four reference genomes to minimize reference bias and reflect information we learned about the sample as we progressed: (1) to produce a phylogeny including six extant Cervidae species, we mapped the data to the cattle, caribou and white-tailed deer nuclear genome references (GCF_002263795.2, GCA_949782905, and GCF_023699985.2, respectively); and (2), because the nuclear genome analyses showed clear *Odocoileus* affinity of the sample, we mapped to the white-tailed deer mitogenome (NC_015247.1) to produce a phylogeny with more species and gain resolution on the systematics. We used bwa aln (v0.7.17)[35] and parameters optimised for aDNA data: a maximum of two gaps (-o 2), allowing more substitution (-n 0.01), and deactivating seed (-l 16500)[36,37]. We filtered mapped reads to a minimum length of 30 bp and a minimum mapping quality of 25 in SAMtools (v1.17)[38]. We identified and removed PCR duplicates with the python script samremovedup.py, which takes into account the read start and end coordinates and read length[39]. We used BamTools stats (v2.5.1)[40] throughout the mapping process for quality checks and computed the final coverage in SAMtools (Table S4). Finally, we assessed damage patterns in mapDamage (v2.2.1)[41] and PMDtools (v2)[42], the latter using the platypus option that allows damage identification at CpG sites which remain unaffected by USER enzyme treatment (Fig. S3).

For the modern cervid resequencing data (Table S3), we downsampled the data to a maximum of 100 million random read pairs using seqtk sample (v1.3)[43] to even out the modern data, making sure the seed was set to the same number for each read pair file. We trimmed the downsampled reads in Trimmomatic (v0.36)[44]. Using bwa-mem (v0.7.17)[35], we mapped the data to the same references as above. We used SAMtools (v1.10)[38] to sort the aligned reads, then used Picard MarkDuplicates (v2.23.2)[45] and Sambamba view (v0.7.0)[46] to identify and filter out duplicated reads. Finally, we used GATK RealignerTargetCreator and IndelRealigner (v4.1.7.0)[47] to carry out a local realignment.

### Divergence within Cervidae

#### Whole genome phylogeny and allelic distance

Because we did not know the species, we first mapped the samples to the Cervidae outgroup (cow) to narrow down species assignment; we created an initial dataset in ANGSD (v0.939)[48] including variant and invariant sites but no missing data to estimate sequence divergence (d_XY_) which we computed using a custom script (popgenWindows.py)[49]. We then called genotypes and genotype likelihoods in ANGSD removing all missing data (-minInd 37) and filtering for a maximum p-value of 1 e−6 (-SNP_pval 1e-6). We further applied filtering using VCFtools (v 0.1.16)[50] and created a minimal linkage disequilibrium filter dataset, where we allowed a maximum of one site per 100 bp (--thin 100; minLD dataset).

We used PLINK (v1.90b6.21)[51] to compute allelic distance (--distance) between all individuals and plotted the resulting matrix in heatmaply[52]. We inferred a time-calibrated Cervidae phylogeny based on SNP data using the multi-species coalescent model in SNAPP (v 1.6.1)[53,54], setting two calibration points following a normal distribution, one on the *Cervus* node (μ = 2.3, σ= 0.5[20,21]) and one on the Odocoileini node (μ = 5.8, σ= 0.5[55]), and a starting tree modifying the allelic distance tree produced above into a species tree. We used a custom script (snapp_prep.rb[54]) with default settings to generate two XML inputs for each dataset: one with all the filtered data and one that included transversions only as they should represent true mutations as opposed to transitions which can be the product of DNA damage (-- transversions). We ran SNAPP in three independent runs of 500,000 MCMC iterations, excluding the first 10% as burn-in. We assessed convergence and checked that the effective sample size was >200 in Tracer (v1.7.2)[56], and generated a maximum clade credibility tree with median node heights using TreeAnnotator (v2.7.6)[57] before visualisation.

As the data mapped to cattle, and indeed all reference genomes (Table S4), suggested a relationship to the *Odocoileus* genus we created the same minLD dataset and repeated the allelic distance and phylogeny analyses with data mapped to caribou and white-tailed deer to see if the placement of ROMM75974 was impacted by reference bias. In addition, we created a minimal LD and depth filter dataset that included a filter for a genotype minimum depth of 3x (--minDP 3; minLD & D3 datasets). We created the latter to increase the confidence in the ancient data, and phylogenetic and allelic distance analyses were performed on both datasets (Table S4).

Finally, because caribou females are antler-bearing, which is unique in Cervidae, identifying the molecular sex of ROMM75974 could inform on phylogenetic affinity. Females have two copies of autosomes and X chromosomes; their X/A ratio is expected to be close to one, whereas it should be approximately 0.5 in males who only have one X copy[58]. We used SAMtools idxstats to retrieve the number of reads mapped to each chromosome of the caribou and cattle reference, using chromosomes one and two, respectively, as they are closest in size to their X chromosome.

#### Mitochondrial analyses

We created a mitochondrial consensus sequence for ROMM75974 in ANGSD, setting a minimum depth of 3 (-setMinDepth 3) and selecting the base with the highest effective depth (-doFasta 3) which resulted in a 10,823 base pair sequence (hereafter referred to as partial mitogenome). We used MEGA (v11.0.13)[59] and the ClustalW algorithm with default settings to align the partial mitogenome sequence to that of 17 other Cervidae species (Table S2), removing all gaps, ambiguous sites, and singletons from the alignment. To perform phylogenetic analysis, we used IQ-TREE (v1.6.12)[60–62] and selected the best substitution model with ModelFinder[61], we obtained branch support from the SH-like approximate likelihood ratio test (-alrt 1000) and used ultrafast bootstrap (-bb 1000) optimised with a hill-climbing nearest neighbour interchange search (-bnni). Because of the contradicting position of ROMM75974 in that phylogeny, we further investigated the possibility of incomplete lineage sorting (ILS) within Odocoileini (*Odocoileus, Mazama, Pudu*, etc.) using the same method as above and 21 published Odocoileini mitogenomes (Table S2), as well as with unpublished mitogenome data of 13 *Odocoileus* individuals[63,64] (Table S3).

## Results

We successfully recovered genomic data from the 11-thousand-year-old sample ROMM75974, that presented signs of damage consistent with ancient DNA (Fig. S3), with endogenous content < 0.5% (Table S4). The average depth and number of variants pre-filtering was highest in data mapped to WTD (0.006x and 121,195 sites), suggesting a better match of reference. The number of variants after strict filtering was drastically reduced (<35 sites) impacting topologies (Table S4, Fig. S4, Fig. S5). We further obtained a coverage of 1.8x for data mapped to the white-tailed deer mitogenome and identified the molecular sex of ROMM75974 as male in all datasets (Table S4).

The nuclear genome phylogenies and allelic distance analyses comprising sufficient informative sites consistently showed ROMM75974 as a sister taxon to the white-tailed – mule deer (*O. hemionus*) clade, with one exception (Fig. S4, Fig. S5). Inferred divergence times ranged between 1.9 and 3 Mya, noting error bars overlapped among *Odocoileus* splits (Fig. 2, Fig. S5). Allelic distance analysis suggested ROMM75974 as closest to both *Odocoileus* species, with an average distance of 22.6% from the *Odocoileus* individuals, which is similar to the WTD – MD distance of 19%, as opposed to 40.5% from all others (Fig. 2, Fig. S4), noting the dxy preliminary analysis support this finding (Table S5). Lastly, ROMM75974 was placed as sister to mule deer in the partial mitogenome phylogeny (Fig. 3).

**Figure 2:**
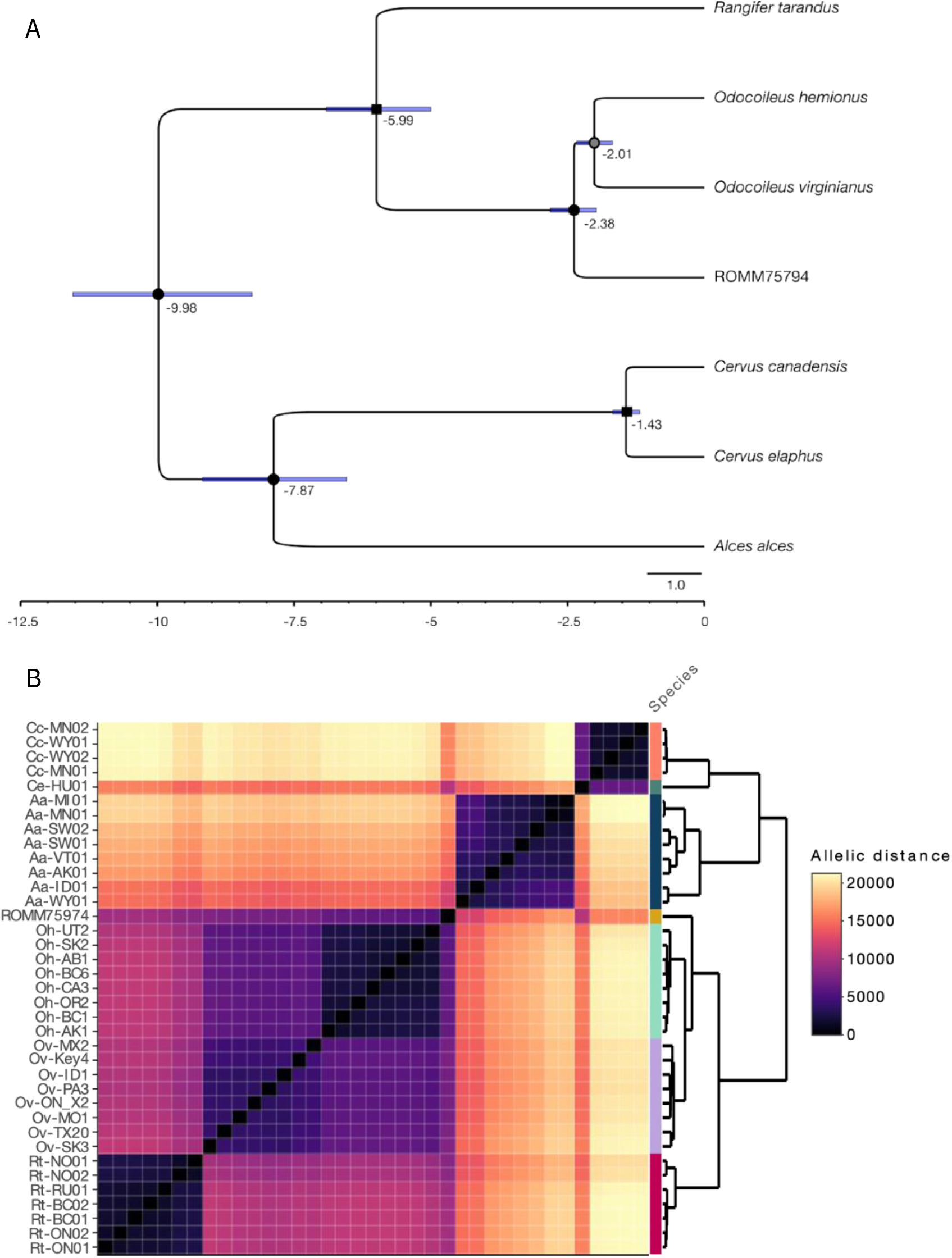
Nuclear analyses on data mapped to WTD. A) Calibrated whole genome phylogeny of Cervidae produced by SNAPP based on the minimal LD filter dataset without transitions (9,255 sites). Nodes are labelled with the estimated divergence time in the past, colour represents posterior distribution support (white ≤ 70%, grey 71 -89%, black ≥ 90%), square nodes show calibration points and blue bars represent the 95% HPD intervals, x-axis timeline in units of millions of years ago (Mya). Tree visualisation performed in FigTree and edited in Inkscape. B) Allelic distance matrix heatmap and clustering based on minimal LD filter dataset (30, 040 sites) of Cervidae, side colour represents the species as abbreviated in sample name: Aa = *Alces alces*, Cc = *Cervus canadensis*, Ce = *Cervus elaphus*, Oh = *Odocoileus hemionus*, Ov = *Odocoileus virginianus*, Rt = *Rangifer tarandus*.

**Figure 3:**
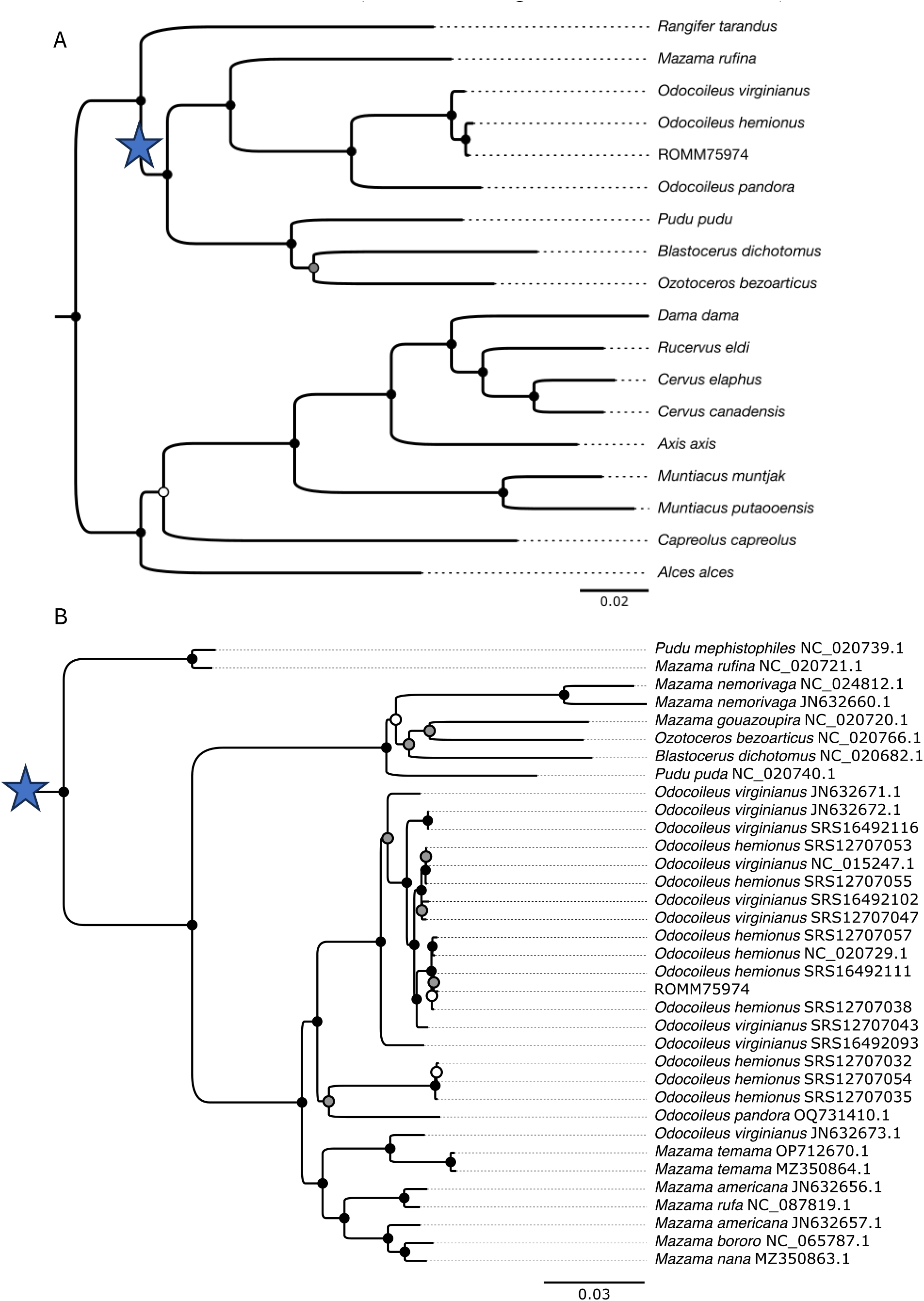
Partial mitogenome phylogeny of Cervidae (A; 6867 sites), and Odocoileini (B; 7018 sites) produced by IQtree. Blue star depicts the Odocoileini branch, accession numbers follow species name in (B). Nodes colour represents bootstrap support (white ≤ 70%, grey 71 - 89%, black ≥ 90%). Tree visualisation performed in FigTree and edited in Inkscape.

## Discussion

Pleistocene extinction studies have focussed on megafaunal mammals at the genus-level because fossil identification to the species-level is often difficult. Genus-level inferences have several shortcomings; they quantitatively minimise the biodiversity loss by not accounting for species numbers[65] (i.e. up to five species in extinct North American genera[6]); they also do not include extinct species whose genera survived (e.g., *Panthera spelaea*[66]). A genus is not representative of functional diversity as species within the same genus can have distinctive functional traits (e.g., *Ursus arctos & Ursus maritimus*) and their loss can have drastically different impacts on the ecosystem[67]. The advent of ancient DNA analyses and museomics allows for reconstructing the late Pleistocene extinction event with more resolution, and to fully quantify the loss of biodiversity that took place at the species and even population level[68].

### Taxonomic resolution of the Subway deer

The rarity of *T. hypogaeus* in the fossil record (only one specimen) and its description based on antler morphology has contributed to the uncertainty around the taxon’s designation as a monotypic genus, with some early studies assigning the specimen to *Rangifer*[22,29]. Our results consistently invalidated ROMM75974 as a member of *Rangifer*; rather, all analyses including sufficient data showed the specimen as closest to *Odocoileus*.

The nuclear genome phylogenies have a general topology of Cervidae that is consistent with [69], and generally suggest ROMM75974 as a sister taxon to the white-tailed – mule deer clade (Fig. 2A, Fig. S5). One nuclear phylogeny does show the specimen as sister to mule deer (Fig. S5), but this pattern disappears with the transversions only filter, suggesting it might be an artefact of ancient DNA. We unfortunately could not recover enough sites at a depth of three on data mapped to WTD and caribou, though 1x coverage produces reliable inferences in phylogenetic and population structure analyses (e.g. [70,71]).

The partial mitogenome phylogeny also showed ROMM75974 as embedded in a mule deer clade (Fig. 3). This result at the mitogenome level is, however, to be taken with some caution as the discrepancy between nuclear and mitochondrial phylogeny is well known in white-tailed and mule deer for which the mito-nuclear discordance has been a long-standing debate (e.g. [72–74]). More broadly, the phylogenetic relationships of the Odocoileini remain largely unresolved[69], and mitochondrial phylogenies of the group often present paraphyly[69,75] (Fig. 3B), with both *Odocoileus* and *Mazama* showing poor species-level resolution. Highlighting this, *O. pandora*, which was recently transferred to *Odocoileus* from *Mazama* [76] was included in the mitochondrial phylogeny, and like *Torontoceros*, is grouped in a paraphyletic fashion . Given the lack of historical nuclear gene flow between white-tailed and mule deer[63], the mito-nuclear discordance we observe here likely results from ILS. The collective evidence, genomic and antler, suggest ROMM75974 was a phenotypically and genetically unique lineage. Additional support for this argument is the close relationship of ROMM75974 to *O. hemionus*; this species is found exclusively in the western half of the continent with no evidence of suitable historical habitat in the east[26,77].

It is clear that the specimen belongs to *Odocoileus*, either as a separate extinct species, or as a member of an extinct lineage that diverged from *hemionus*. Given that two phylogenies shows ROMM75974 as sister to mule deer, we are unable to differentiate these two scenarios with certainty. Still, we suggest that this pattern is the result of rapid divergence, ILS and ancient DNA artefacts, as most phylogenies support species divergence (Fig. S5). If ROMM75974 were a representative of a different species, it would have been part of a rapid *Odocoileus* radiation event that took place 1-3 Mya from a common ancestor, which resulted in limited resolution of the mitogenome phylogeny across Odocoileini. Recent gene flow between white-tailed and mule deer could bring modern representatives of those species closer together in phylogenies and sequence divergence compared to Pleistocene individuals. Comparing ROMM75974 to Pleistocene representatives of white-tailed and mule deer and using additional and high-quality aDNA analyses would help confirm a distinct species designation.

If the range of the Subway deer was restricted to the North American Great Lakes, finding additional members of the taxa will likely be challenging as the fossil record in the region is limited for Cervidae[78]; it is possible that it ranged further south and west if the specimen represents a divergent, but extinct mule deer lineage. Nevertheless, an integrative taxonomy approach to the species identification of a museum specimen incorporating genomic data is increasingly common practice[79–83], and arguably more reliable than morphological assignment, particularly based on antler structure[14]. As fossil identification to the species level of this genus is difficult[15,84], DNA assessment of additional *Odocoileus* museum specimens might identify other representatives of this potential new species and give a better idea of genetic variation prior to extinction (e.g. [85,86]).

### Adding to the Ice Age extinction list of North America

The late Pleistocene extinction event was global and affected many species. While megafaunal mammals received most of the attention, particularly the emblematic Proboscidea (*Mammut, Mammuthus*), smaller mammals, birds and trees were also impacted[13,87,88]. North America saw over 30 mammal genera disappear from the continent[1,6,9,12].

The late Pleistocene landscape of southern Ontario, the location of current day Toronto, and more broadly the North American Great Lakes region, was similar to boreal woodland, dominated by spruce and sedge[89–92]. The area was home to a variety of extant fauna such as caribou and grizzly bears (*Ursus arctos–horribilis*)[92–95] whose ranges have significantly shifted, but also the extinct mammoth (*Mammuthus sp*.)[7,92,96–98], mastodon (*Mammut americanum*)[97,98] and stag-moose (*C. scotti*)[93,99,100]. One explanation for these range shifts and extinctions is the rapid vegetal transition from an open boreal woodland habitat to a more closed pine forest by the end of the Younger Dryas[7,77,89,90,93,98,101,102]. If ROMM75974 represented a distinct but extinct *Odocoileus* species or population, it would have been strongly impacted by this sudden vegetation replacement, as Croitor (2022)[22] suggested its antler morphology reflected an adaptation to open landscapes. While our results exclude the genus *Torontoceros* from future assessments of the Late Pleistocene extinction event, the aDNA assessment support it being included as an extinct *Odocoileus* species or lineage. Doing so would help highlight the role or climate change in some megafaunal extinctions, although their causes are highly variable and species-specific.

## Acknowledgements

The specimen ROMM75974 was discovered on the ancestral land of different First Nations including the Anishinaabeg, the Haudenosaunee, and the Wendat. Historically, the region of Toronto was a busy trade route which connected many First Nations of the Great Lakes, it was a place of culture, important for fishing and hunting, and generally highly significant to the people who lived there for thousands of years. The land on which we work belongs to many First Nations, it is their ancestral home and unceded territory. Together, Trent University and the ROM are located on the territory of the Michi Saagiig Anishnaabeg, the Wendat, the Haudenosaunee Confederacy, and the Anishinaabeg Nation, including the Mississaugas of the Credit First Nation. As settlers, we are grateful to have had the opportunity to live and work on this land, and to benefit from it. We would like to show our respect to the First Peoples and thank them for their care, stewardship, and teachings. Miigwetch, tiawenhk, niá:wen.

Camille Kessler was supported by an International Graduate Scholarship, an Ontario Graduate Scholarship, and a French American Charitable Trust Scholarship. This work was supported by NSERC Discovery Grant (Grant Number: RGPIN-2017-03934); ComputeCanada Resources for Research Groups (Grant Number: RRG gme-665-ab); Canadian Foundation for Innovation: John R. Evans Leaders Fund and the Ontario Early Researcher Award (Grant Number: #36905). We would like to thank Marianne Dehasque for their precious help and sharing of laboratory methods and protocols, Marie-Laurence Cossette for their help in the lab, and Jochen Wolf and Jose Alberto Lopez-Aleman for helpful comments.

