## Supplementary figures and tables for "Ancient DNA of the Toronto Subway Deer Adds to the Extinction List of Ice Age Megafauna"

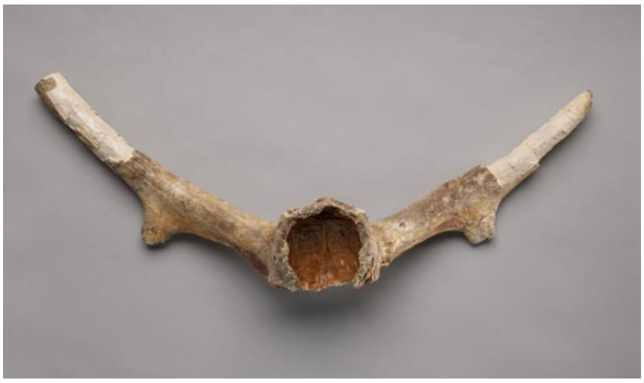
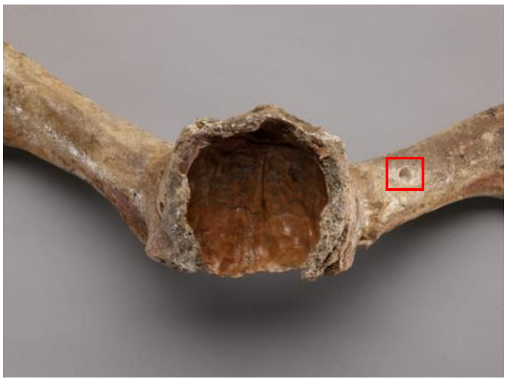
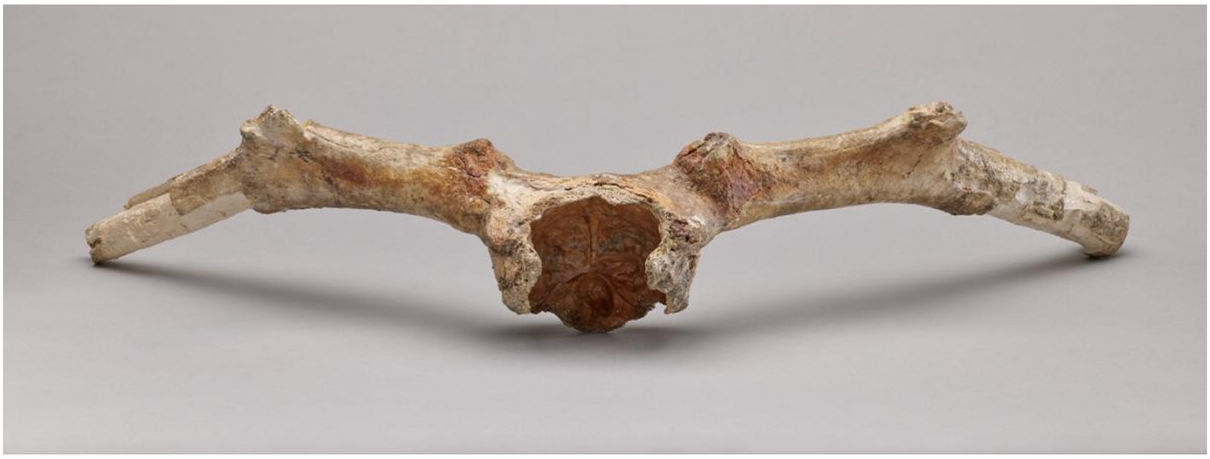
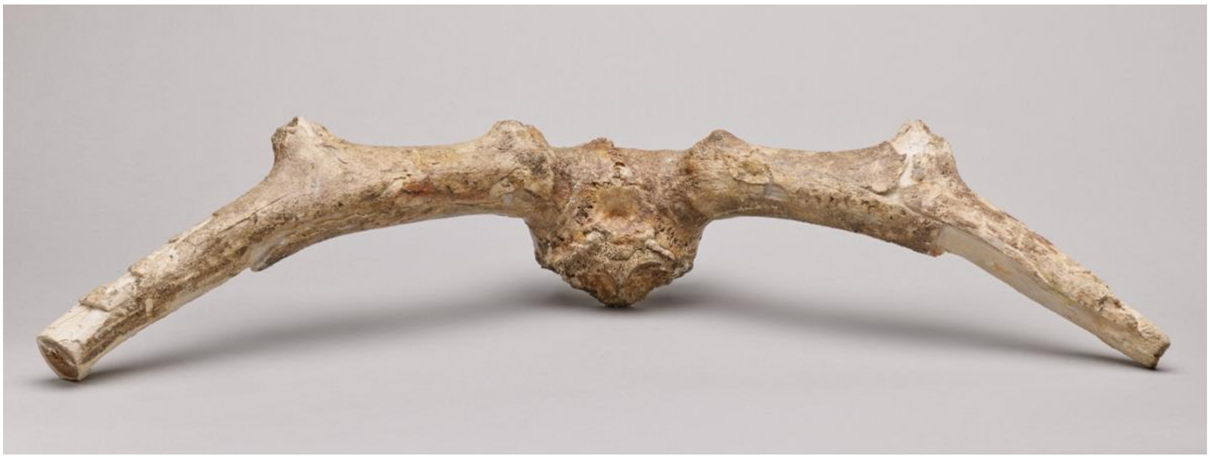
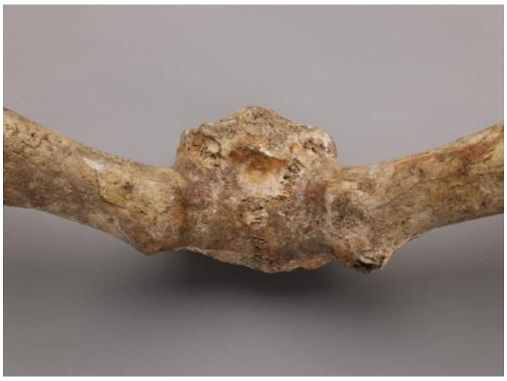

Figure S1: Specimen ROMM75974 from the Royal Ontario Museum, Toronto, additional pictures, red square indicates sampling location.

Figure S2: Approximate sampling locations, coloured by species.

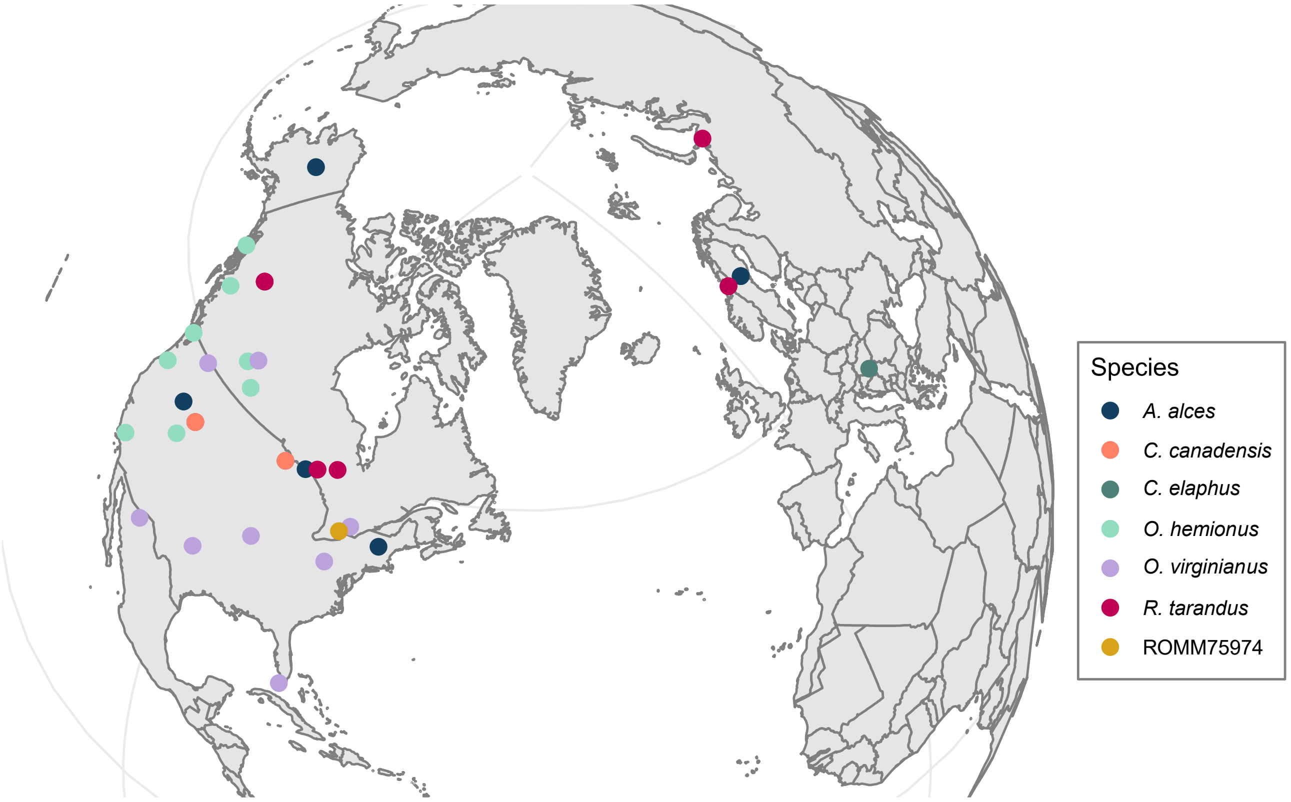

Figure S3: ROMM75974 damage patterns from MapDamage (A) and PMDtools (B) when mapped to caribou. Deamination at CpG sites identified by PDMtools is unaffected by USER enzyme treatment.

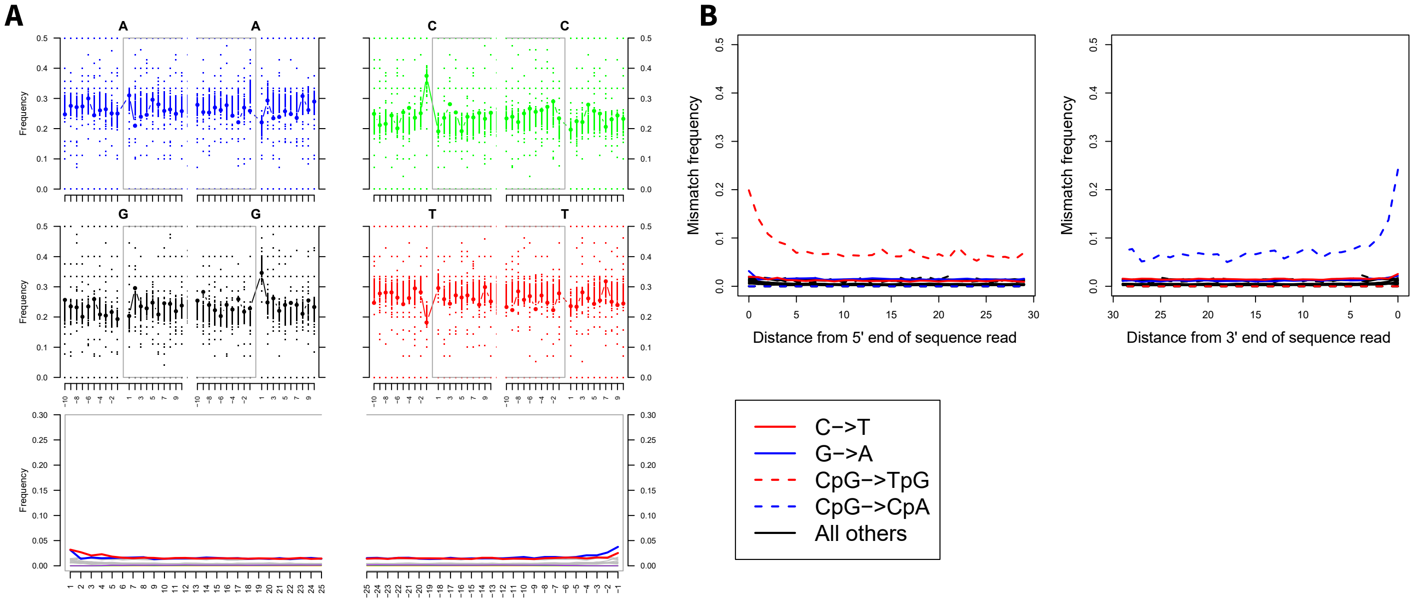

Figure S4: Allelic distance matrix heatmaps and clustering based on (A) WTD minLD dataset (30,040 sites), (B) WTD minLD & D3 dataset (32 sites), (C) caribou minLD dataset (29,095 sites), (D) caribou minLD & D3 dataset (16 sites), and (E) cattle minLD dataset (exclusively part of initial screening, 11,882 sites). Side colour represents the species as abbreviated in sample name: Aa = *Alces alces*, Cc = *Cervus canadensis*, Ce = *Cervus elaphus*, Oh = *Odocoileus hemionus*, Ov = *Odocoileus virginianus*, Rt = *Rangifer tarandus*, yellow star indicates non-consistent topology.

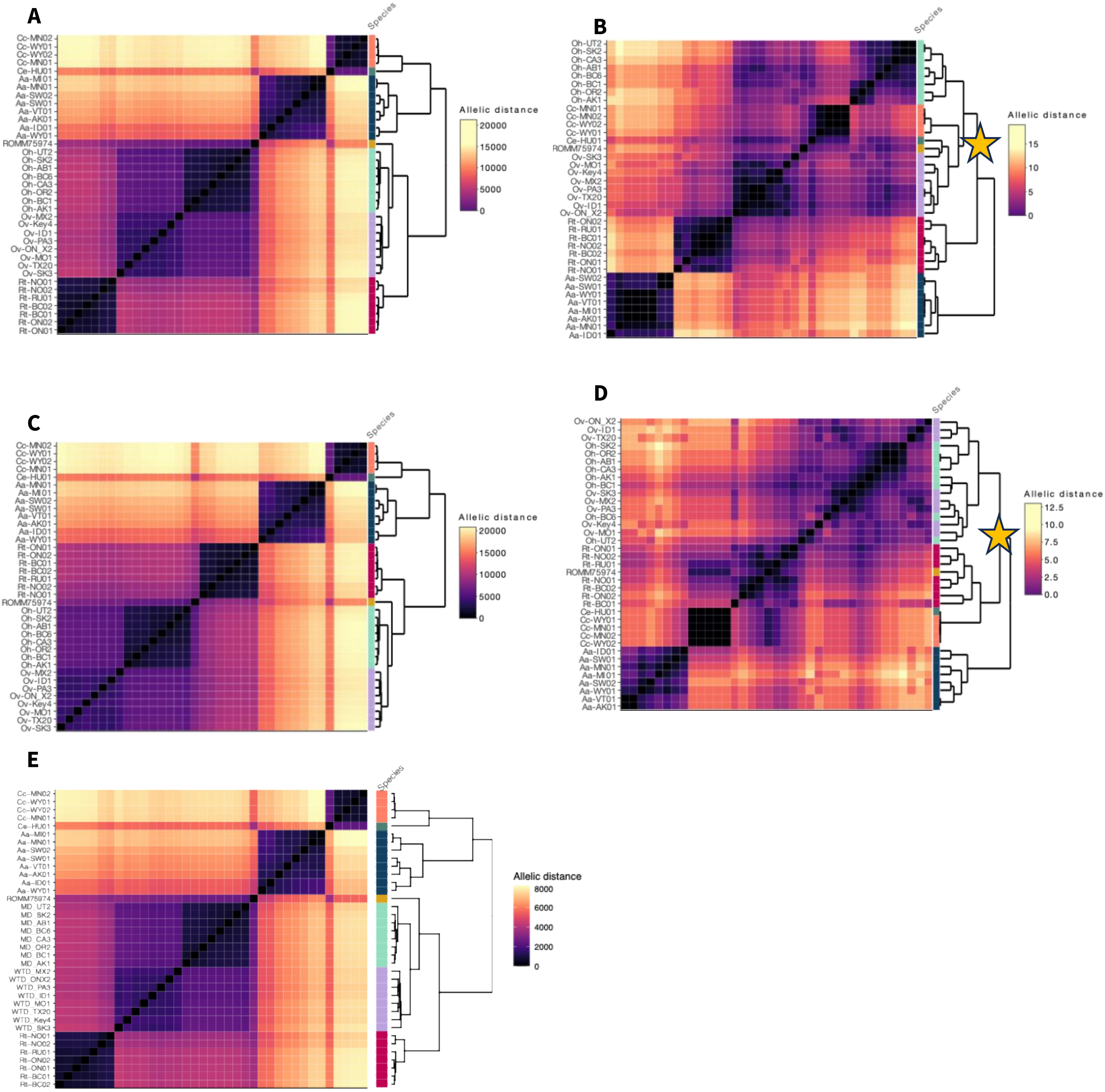

Figure S5: Calibrated whole genome phylogeny of Cervidae produced by SNAPP, nodes are labelled with the estimated divergence time in the past, node colour represents posterior distribution support (white ≤ 70%, grey 71 - 89%, black ≥ 90%), square nodes show square nodes show calibration points and blue bars represent the 95% HPD intervals, a yellow star indicates non-consistent topology, and red number of sites means combined EES < 200 for the likelihood, prior and posterior distribution. Cattle phylogenies as part of initial screening, transv datasets mean transversions only.

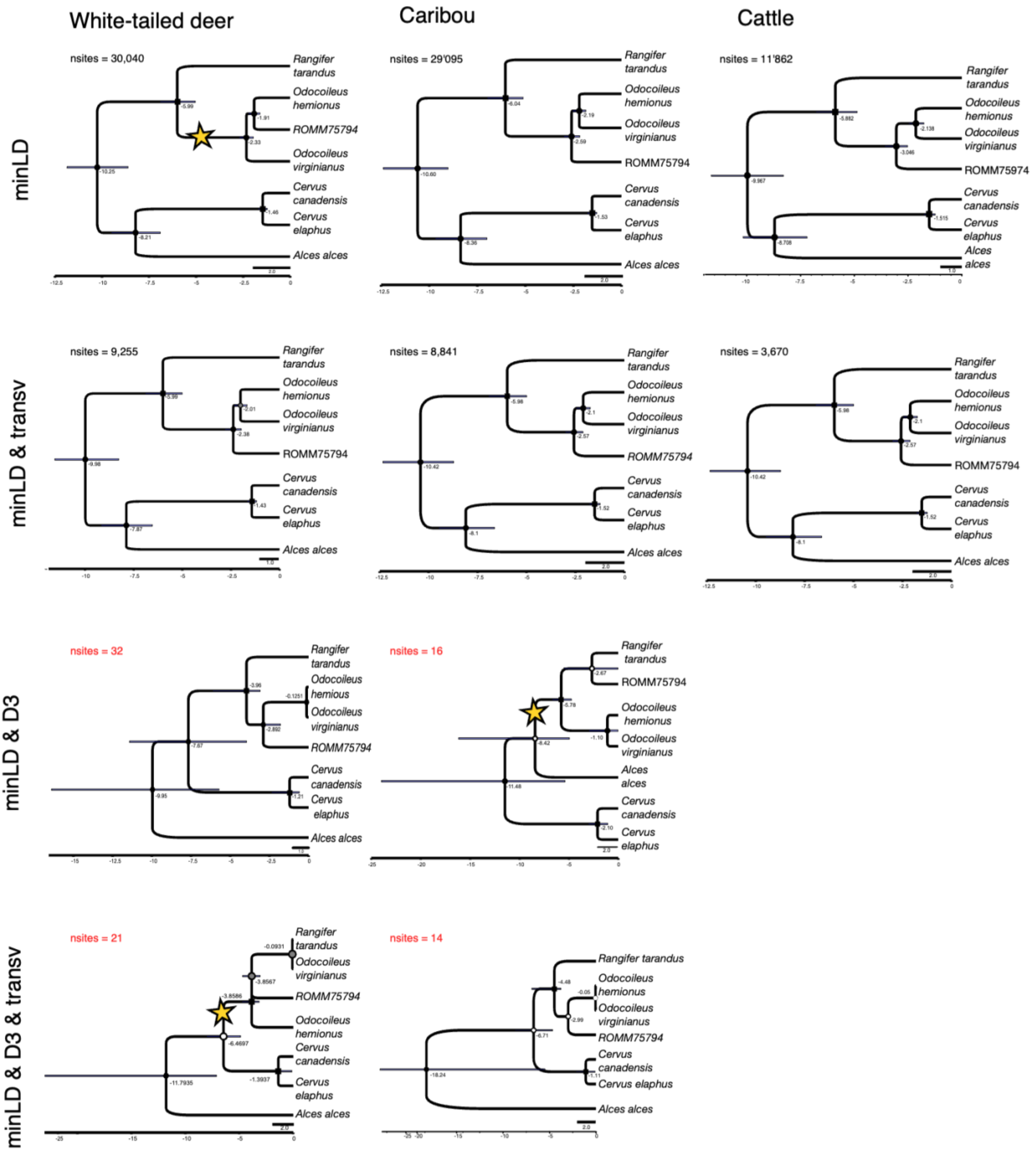

**Supplementary tables**

Table S1: Mini literature review of papers investigating extinction and including or excluding *Torontoceros hypogeaus*.

| List of extinct NA mammals | Examples |
|---|---|
| Included | Barnosky, A.D., Koch, P.L., Feranec, R.S., Wing, S.L., and Shabel, A.B. (2004). Assessing the Causes of Late Pleistocene Extinctions on the Continents. Science 306, 70–75. 10.1126/science.1101476.<br>Elias, S., and Schreve, D. (2007). Late Pleistocene Megafaunal Extinctions. In Encyclopedia of Quaternary Science. (Elsevier), pp. 3202–3217. 10.1016/B0-444-52747-8/00266-0.<br>Jackson, L.J. (1988). Fossil Cervids and Fluted Point Hunters: A Review for Southern Ontario. Ontario Archaeology OA48, 15.<br>Koch, P.L., and Barnosky, A.D. (2006). Late Quaternary Extinctions: State of the Debate. Annu. Rev. Ecol. Evol. Syst. 37, 215–250. 10.1146/annurev.ecolsys.34.011802.132415.<br>Martin, P.S., and Steadman, D.W. (1999). Prehistoric Extinctions on Islands and Continents. In Extinctions in Near Time, R. D. E. MacPhee, ed. (Springer US), pp. 17–55. 10.1007/978-1-4757-5202-1_2.<br>Meltzer, D. J., & Mead, J. I. (1985). Dating late Pleistocene extinctions: theoretical issues, analytical bias, and substantive results. Environments and Extinctions: Man in Late Glacial North America, 145-173.<br>Spaulding, W.G. (1983). The Overkill Hypothesis as a Plausible Explanation for the Extinctions of Late Wisconsin Megafauna. Quat. res. 20, 110–112. 10.1016/0033-5894(83)90069-8.<br>Faith, J.T., and Surovell, T.A. (2009). Synchronous extinction of North America's Pleistocene mammals. Proc. Natl. Acad. Sci. U.S.A. 106, 20641–20645. 10.1073/pnas.0908153106. |
| Excluded | Grayson, D.K. (2007). Deciphering North American Pleistocene Extinctions. Journal of Anthropological Research 63, 185–213. 10.3998/jar.0521004.0063.205.<br>Grayson, D.K. (1991). Late Pleistocene mammalian extinctions in North America: Taxonomy, chronology, and explanations. J World Prehist 5, 193–231. 10.1007/BF00974990.<br>Meltzer, D.J. (2020). Overkill, glacial history, and the extinction of North America's Ice Age megafauna. Proc. Natl. Acad. Sci. U.S.A. 117, 28555–28563. 10.1073/pnas.2015032117.<br>Stewart, M., Carleton, W.C., and Groucutt, H.S. (2021). Climate change, not human population growth, correlates with Late Quaternary megafauna declines in North America. Nat Commun 12, 965. 10.1038/s41467-021-21201-8.<br>Stuart, A.J. (2015). Late Quaternary megafaunal extinctions on the continents: a short review. Geol. J. 50, 338–363. 10.1002/gj.2633. |

Table S2: Information regarding species and mitochondrial genomes used in the phylogenetic analyses of Cervidae and/or Odocoileini. Fossil record age ranges from the The Paleobiology Database [1].

| Species information | | Mitochondrial genome information | | |
|---|---|---|---|---|
| Species | Fossil record age range | Description | Accession | Phylogeny |
| North american species | | | | |
| *Alces Alces* | 2.58 – 0 Ma | Alces alces isolate Kazakhstan mitochondrion, complete genome | NC_020677.1 | Cervidae |
| *Cervus canadensis* | 0.77 – 0 Ma | Cervus canadensis isolate 19FC017 mitochondrion, complete genome | NC_050863.1 | Cervidae |
| *Odocoileus hemionus* | 0.77 – 0 Ma | Odocoileus hemionus isolate T1766 mitochondrion, complete genome | NC_020729.1 | Cervidae & Odocoileini |
| *Odocoileus virginianus* | 3.6 – 0 Ma | Odocoileus virginianus mitochondrion, complete genome | NC_015247.1 | Cervidae & Odocoileini |
| | | Odocoileus virginianus isolate CYTO mitochondrion complete genome | JN632672.1 | Odocoileini |
| | | Odocoileus virginianus isolate MRGOv14 mitochondrion complete genome | JN632673.1 | Odocoileini |
| | | Odocoileus virginianus isolate T4887 mitochondrion complete genome | JN632671.1 | Odocoileini |
| *Rangifer tarandus* | 2.58 – 0 Ma | Rangifer tarandus mitochondrion, complete genome | NC_007703.1 | Cervidae |
| Central american species | | | | |
| *Mazama temama* | NA | Mazama temama voucher MZFC:4668 mitochondrion complete genome | OP712670.1 | Odocoileini |
| | | Mazama temama voucher NUPECCE:T366 mitochondrion complete genome | MZ350864.1 | Odocoileini |
| *Odocoileus pandora* | NA | Odocoileus pandora voucher NUPECCE T365 topotype mitochondrion, complete genome | OQ731410.1 | Cervidae & Odocoileini |
| South american species | | | | |
| *Blastocerus dichotomus* | 2.58 – 0 Ma | Blastocerus dichotomus isolate MRGBd8 mitochondrion, complete genome | NC_020682.1 | Cervidae & Odocoileini |
| *Mazama americana* | 0.13 – 0 Ma | Mazama americana isolate MAZ9472 mitochondrion complete genome | JN632656.1 | Odocoileini |
| | | Mazama americana isolate MRGMa40 mitochondrion complete genome | JN632657.1 | Odocoileini |
| *Mazama bororo* | NA | Mazama bororo voucher NUPECCE T215 mitochondrion complete genome | NC 065787.1 | Odocoileini |
| *Mazama gouazoupira* | NA | Mazama gouazoupira isolate MRGsp2 mitochondrion complete genome | NC 020720.1 | Odocoileini |

| | | | | |
|---|---|---|---|---|
| *Mazama nana* | NA | Mazama nana voucher NUPECCE:T107 mitochondrion complete genome | MZ350863.1 | Odocoileini |
| *Mazama nemorivaga* | NA | Mazama nemorivaga isolate MRGMa36 mitochondrion complete genome | NC 024812.1 | Odocoileini |
| | | Mazama nemorivaga isolate T1627 mitochondrion complete genome | JN632660.1 | Odocoileini |
| *Mazama rufa* | NA | Mazama rufa mitochondrion complete genome | NC 087819.1 | Odocoileini |
| *Mazama rufina* | NA | Mazama rufina isolate MRGMr4 mitochondrion, complete genome | NC_020721.1 | Cervidae & Odocoileini |
| *Ozotoceros bezoarticus* | 0.13 – 0 Ma | Ozotoceros bezoarticus isolate MRGOb2 mitochondrion, complete genome | NC_020766.1 | Cervidae & Odocoileini |
| *Pudu mephistophiles* | NA | Pudu mephistophiles isolate MRGPm2 mitochondrion complete genome | NC 020739.1 | Odocoileini |
| *Pudu puda* | NA | Pudu puda isolate M92144 mitochondrion, complete genome | NC_020740.1 | Cervidae & Odocoileini |
| Eurasian species | | | | |
| *Axis axis* | 0.77 – 0 Ma | Axis axis isolate CYTO mitochondrion, complete genome | NC_020680.1 | Cervidae |
| *Capreolus capreolus* | 2.58 – 0 Ma | Capreolus capreolus isolate CYTO mitochondrion, complete genome | NC_020684.1 | Cervidae |
| *Cervus elaphus* | 3.6 – 0 Ma | Cervus elaphus mitochondrion, complete genome | NC_007704.2 | Cervidae |
| *Dama dama* | 2.58 – 0 Ma | Dama dama isolate CYTO mitochondrion, complete genome | NC_020700.1 | Cervidae |
| *Muntiacus muntjak* | 0.13 – 0 Ma | Muntiacus muntjak mitochondrion, complete genome | NC_004563.1 | Cervidae |
| *Muntiacus putaoensis* | NA | Muntiacus putaoensis mitochondrion, complete genome | NC_036430.1 | Cervidae |
| *Rucervus eldii* | 0.77 – 0 Ma | Rucervus eldi mitochondrion, complete genome | NC_014701.1 | Cervidae |

1. 2025 The Paleobiology Database. *The Paleobiology Database*. See https://paleobiodb.org/#/ (accessed on 3 September 2025).

Table S3: Whole genome data summary: SRA accessions and sample information of the 36 modern samples used, coloured by species.

| Species | Sample ID | Accession | Locality | Phylogeny |
|---|---|---|---|---|
| *Alces alces* ● | Aa-AK01 | SRR6079187 | Alaska | Cervidae |
| *Alces alces* ● | Aa-ID01 | SRR6079177 | Idaho | Cervidae |
| *Alces alces* ● | Aa-MI01 | SRR18899225 | Michigan | Cervidae |
| *Alces alces* ● | Aa-MN01 | SRR18899222 | Minnesota | Cervidae |
| *Alces alces* ● | Aa-SW01 | ERR12087977 | Sweden | Cervidae |
| *Alces alces* ● | Aa-SW02 | ERR12087904 | Sweden | Cervidae |
| *Alces alces* ● | Aa-VT01 | SRR6079181 | Vermont | Cervidae |
| *Alces alces* ● | Aa-WY01 | SRR6079200 | Wyoming | Cervidae |
| *Cervus canadensis* ● | Cc-WY01 | SRR12450513 | Wyoming | Cervidae |
| *Cervus canadensis* ● | Cc-MN01 | SRR12450505 | Minnesota | Cervidae |
| *Cervus canadensis* ● | Cc-MN02 | SRR12450502 | Minnesota | Cervidae |
| *Cervus canadensis* ● | Cc-WY02 | SRR12450506 | Wyoming | Cervidae |
| *Cervus elaphus* ● | Ce-HU01 | SRR956941 | Hungary | Cervidae |
| *Rangifer tarandus* ● | Rt-BC01 | SRR27590283 | British Columbia | Cervidae |
| *Rangifer tarandus* ● | Rt-BC02 | ERR11471728 | British Columbia | Cervidae |
| *Rangifer tarandus* ● | Rt-ON01 | ERR11471702 | Ontario | Cervidae |
| *Rangifer tarandus* ● | Rt-ON02 | SRR24951207 | Ontario | Cervidae |
| *Rangifer tarandus* ● | Rt-NO01 | SRR24951210 | Norway | Cervidae |
| *Rangifer tarandus* ● | Rt-NO02 | SRR15459420 | Norway | Cervidae |
| *Rangifer tarandus* ● | Rt-RU01 | SRR15459424 | Russia | Cervidae |
| *Odocoileus hemionus* ● | Oh_AB1 | SRS12707053 | Alberta | Cervidae & Odocoileini |
| *Odocoileus hemionus* ● | Oh_AK1 | SRS12707054 | Alaska | Cervidae & Odocoileini |
| *Odocoileus hemionus* ● | Oh_BC1 | SRS12707032 | British Columbia | Cervidae & Odocoileini |
| *Odocoileus hemionus* ● | Oh_BC6 | SRS12707055 | British Columbia | Cervidae & Odocoileini |
| *Odocoileus hemionus* ● | Oh_CA3 | SRS12707057 | California | Cervidae & Odocoileini |
| *Odocoileus hemionus* ● | Oh_OR2 | SRS12707035 | Oregon | Cervidae & Odocoileini |

| | | | | |
|---|---|---|---|---|
| *Odocoileus hemionus* ● | Oh_SK2 | SRS16492111 | Saskatchewan | Cervidae & Odocoileini |
| *Odocoileus hemionus* ● | Oh_UT2 | SRS12707038 | Utah | Cervidae & Odocoileini |
| *Odocoileus virginianus* ● | Ov_ID1 | SRS16492129 | Idaho | Cervidae & Odocoileini |
| *Odocoileus virginianus* ● | Ov_Key4 | SRS16492093 | Florida | Cervidae & Odocoileini |
| *Odocoileus virginianus* ● | Ov_MO1 | SRS16492102 | Missouri | Cervidae & Odocoileini |
| *Odocoileus virginianus* ● | Ov_MX2 | SRS12707043 | Mexico | Cervidae & Odocoileini |
| *Odocoileus virginianus* ● | Ov_ONX2 | SRS16492116 | Ontario | Cervidae & Odocoileini |
| *Odocoileus virginianus* ● | Ov_PA3 | SRS12707047 | Pennsylvania | Cervidae & Odocoileini |
| *Odocoileus virginianus* ● | Ov_SK3 | SRS12707051 | Saskatchewan | Cervidae |
| *Odocoileus virginianus* ● | Ov_TX20 | SRS16492126 | Texas | Cervidae |

Table S4: Quality check for ROMM75974 data mapped to cattle, white-tailed deer and caribou. Final read number represents reads over 25 bp with a minimum quality of 30 and with duplicates removed.

| | Reference | Reads total | Reads mapped (#) | Reads mapped (%) | Final filtered reads (#) | Final average read length | X/A ratio | Depth | called variant # | variant # minLD dataset | variant # minLD & minD dataset |
|---|---|---|---|---|---|---|---|---|---|---|---|
| ROMM75794 | Cattle (GCF_002263795.2) | 26579834 | 61310 | 0.2307 | 39941 | 62 | 0.624 | 0.00005 | 41794 | 11862 | 8551 |
| | Caribou (GCA_949782905.1) | 26579834 | 128407 | 0.4831 | 81022 | 70 | 0.668 | 0.0001 | 107109 | 29095 | 21 |
| | WTD (GCF_023699985.2) | 26579834 | 132615 | 0.4989 | 82435 | 71 | 0.528 | 0.0059 | 121195 | 30040 | 32 |
| | White-tailed deer mtDNA (NC_015247.1) | 26579834 | 252 | - | 237 | 80 | - | 1.796 | - | - | - |

Table 5: Sequence divergence ($d_{XY}$) for data mapped to cattle, used as part of initial screening.

| | | ROMM75974 | *A. alces* | *C. canadensis* | *C. elaphus* | *O. hemionus* | *O. virginianus* | *R. tarandus* |
|---|---|---|---|---|---|---|---|---|
| $d_{XY}$ | ROMM75974 | - | 0.0170 | 0.0183 | 0.0144 | 0.0089 | 0.0106 | 0.0122 |
| | *A. alces* | 0.0170 | - | 0.0232 | 0.0193 | 0.0198 | 0.0200 | 0.0206 |
| | *C. canadensis* | 0.0183 | 0.0232 | - | 0.0088 | 0.0233 | 0.0236 | 0.0241 |
| | *C. elaphus* | 0.0144 | 0.0193 | 0.0088 | - | 0.0185 | 0.0188 | 0.0194 |
| | *O. hemionus* | 0.0089 | 0.0198 | 0.0233 | 0.0185 | - | 0.0092 | 0.0138 |
| | *O. virginianus* | 0.0106 | 0.0200 | 0.0236 | 0.0188 | 0.0092 | - | 0.0139 |
| | *R. tarandus* | 0.0122 | 0.0206 | 0.0241 | 0.0194 | 0.0138 | 0.0139 | - |
